# Structural and functional analyses of viral H2 protein of the vaccinia virus entry fusion complex

**DOI:** 10.1101/2023.08.28.555102

**Authors:** Chi-Fei Kao, Chang-Yi Liu, Chia-Lin Hsieh, Kathleen Joyce D. Carillo, Der-Lii M. Tzou, Hao-Ching Wang, Wen Chang

**Affiliations:** Institute of Molecular Biology, Academia Sinica, Taipei, Taiwan; The PhD Program for Translational Medicine, College of Medical Science and Technology, Taipei Medical University and Academia Sinica, Taipei, Taiwan; Graduate Institute of Translational Medicine, College of Medical Science and Technology, Taipei Medical University, Taipei 110, Taiwan; Institute of Chemistry, Academia Sinica, Taipei, Taiwan

**Keywords:** Vaccinia virus, Entry fusion complex, H2 protein crystallization

## Abstract

Virus-mediated membrane fusion involves conformational changes of the viral fusion protein to fuse the opposing viral and host lipid bilayers. Unlike all other known viruses that contain a single fusion protein, poxviruses harbor a multimeric protein complex of 11 subunits, termed the entry fusion complex (EFC), to mediate fusion with host membranes. Yet, how the poxviral EFC mediates membrane fusion remains enigmatic. To establish the mechanism of EFC-triggered membrane fusion, we are deciphering the structure and function of individual EFC components. Here, we determined the crystal structure of the H2 ectodomain by X-ray diffraction, revealing a folded conformation comprising a central five-stranded β-sheet and three cladding α-helices. We reconstructed the full-length H2 by *in silico* prediction, revealing that the N-terminal region (aa 51-90) of H2 protein may fold as a long helix connecting the ectodomain and transmembrane region. Using alanine-mutagenesis screening in a transient complementation system, coimmunoprecipitation, isothermal titration calorimetry and MV-triggered membrane fusion assays, we concluded that the surface of the ectodomain of H2 protein, including two loop regions, ^170^LGYSG^174^ and ^125^RRGTGDAW^132^, constitutes a broad A28-binding region. Moreover, although not involved in A28 binding, the N-terminal helical region approximal to the transmembrane part, encompassing ^64^RIK^66^, ^72^W, and ^83^ESDRGR^88^, is also crucial for viral EFC formation and MV infectivity.

## Introduction

Virus-cell fusion, driven by the viral fusion protein, is essential for enveloped viruses to enter host cells. Upon binding to cellular receptors, a viral fusion protein on the virion surface initiates a sequence of conformational changes to insert its hydrophobic fusion loop or fusion peptide into the host membrane to activate hemifusion and ultimately complete host and viral membrane fusion to release the viral genome into cells [1, 2]. Factors such as an acidic environment, calcium cations, and enzymatic cleavage are known to promote viral membrane fusion [3, 4]. The mechanistic process of viral membrane fusion has become an attractive target for rational antiviral and vaccine design [5].

Vaccinia virus is the prototypic member of the poxviridae [6–8]. Cell entry of poxviruses requires two sets of envelope proteins responsible for attachment and membrane fusion, respectively [9]. The former consists of D8 [10], H3 [11], A27 [12] and A26 [13] that bind to cellular glycosaminoglycans or laminins, whereas the latter engages 11 transmembrane proteins—A16 [14], A21 [15], A28 [16, 17], F9 [18], G3 [19], G9 [20], H2 [21], J5 [22], L1 [23], L5 [24] and O3 [25]—to form the entry fusion complex (EFC) that drives membrane fusion with host lipid bilayers [26, 27]. These previous studies of EFC components have shown that the EFC is solely devoted to virus membrane fusion and that repressing the expression of individual EFC components does not interfere with viral gene expression or virion morphogenesis. Instead, mature virus particles produced by mutant viruses deficient in EFC protein components still attach to cells but are defective at various stages of the membrane fusion process [28], with the bipolar distribution of the EFC on virion surfaces also being disrupted [29]. In the previous studies, three subcomplexes— namely A16:G9 [30, 31], L5:G3 [32], and A28:H2 [33]—have been reported to physically interact with each other despite the absence of other EFC components. Among them, the A16:G9 subcomplex participates in regulating membrane fusion by interacting with the fusion suppressors A26 [34, 35] or the A56/K2 complex in an acid-dependent manner [35, 36]. Interestingly, evolutionary studies have uncovered that not only poxviruses but also many nucleocytoplasmic large DNA viruses (NCLDV) also host viral genes encoding homologs of some EFC components, implying the presence of a conserved membrane fusion mechanism mediated by these DNA viruses [37].

Despite these studies, the biochemical nature of individual EFC components and the molecular mechanism of EFC-triggered membrane fusion remain poorly understood. Although the crystal structures of the A16/G9 [38] and G3/L5 [39] subcomplexes have recently been elucidated, a complete functional characterization of the viral EFC is still lacking. The unique and complex organization of the vaccinia EFC may underscore an unidentified structural class of viral fusion proteins. Establishing the fusion mechanism employed by vaccinia would provide novel insights into the evolution of the virus-cell fusion process.

H2 is a type II transmembrane protein interacting with A28 to form an EFC subcomplex [21]. In vaccinia virus, H2 consists of 189 amino acids (aa), with four cysteines that produce two intramolecular disulfide bridges. A focused study on its highly conserved C-terminal region (aa162–182) revealed an “LGYSG” sequence crucial for vaccinia virus infectivity and interaction of H2 with A28 [40]. However, the peptide subjected to functional analyses in that study was minimal and the authors did not provide any structural insights into H2 loss of function.

Here, we provide a comprehensive, structure-function characterization of H2 by resolving the crystal structure of the H2 ectodomain and by identifying critical H2 residues that contribute to viral infectivity, A28 binding, and viral fusion activity. We also used a state-of-the-art structural prediction tool to investigate a distinct membrane-proximal region adjacent to the transmembrane helix. Hence, we obtain an overall understanding of the vaccinia H2 structure essential for its function.

## Materials and Methods

### Cell cultures and viruses

Human embryonic kidney (HEK) 293T and green monkey kidney BSC40 cells were maintained in complete growth media comprising 1X Dulbecco’s modified Eagle’s medium (DMEM) supplemented with 10% fetal bovine serum (FBS), 100 units/mL of penicillin and 100 µg/mL of streptomycin (Invitrogen). The H2-inducible virus vH2i, a kind gift from Bernard Moss [21], was prepared in BSC40 cells in the presence of 50 µM isopropyl-β-d-thiogalactopyranoside (IPTG).

### Construction of H2 expression plasmid and site-directed mutagenesis

To produce recombinant vaccinia H2 (H2R) protein in *Escherichia coli*, the coding region of the truncated ectodomain of H2 protein (tH2) containing amino acids 91-189 was cloned into an expression system (smt3/Ulp) provided by Dr. C. D. Lima for protein expression and purification, as previously described [41]. The resulting pET28a-Smt3-tH2 construct expresses vaccinia tH2 protein with a10xHis-SUMO tag at its N-terminal region. All subsequent tH2 mutants expressed in bacteria were derived from pET28a-Smt3-tH2 using a QuikChange Lightning Site-Directed Mutagenesis Kit (Agilent) following the manufacturer’s instructions, with mutation accuracy confirmed by sequencing (Genomics Inc., Taiwan).

To construct an expression vector for H2 protein expression in mammalian cells, the vaccinia H2R gene sequences were codon-optimized and cloned into a mammalian expression vector pEF6, denoted pEF6-H2R. All the H2 mutant constructs were subsequently derived from the pEF6-H2R plasmid using a QuikChange Lightning Site-Directed Mutagenesis Kit (Agilent), with mutation accuracy confirmed by sequencing (Genomics Inc., Taiwan).

### Expression and purification of the truncated H2 ectodomain (tH2) in *E. coli*

Wild type or mutant pET28a-Smt3-tH2 plasmid was transformed into SHuffle^®^ T7 Express *lysY* Competent *E. coli* and incubated with LB broth containing 50 μg/ml kanamycin at 30 °C until the optical density (OD_600_) reached 0.4-0.6. Then, recombinant protein expression was induced using 0.4 mM IPTG at 16 °C for 20 h. The resulting bacterial pellets were resuspended in lysis buffer (20 mM Tris-Base pH 8.0, 250 mM NaCl, 5% glycerol and 20 mM imidazole) and lysed using a Low Temperature Ultra-High-Pressure Cell Crusher (JNBIO) at 1600 bar. Initial purification of His-SUMO-tH2 was performed by HisTrap™ excel (5 ml) affinity chromatography (Cytiva). The His-SUMO-tH2 was eluted using an imidazole gradient from 20 mM to 500 mM (binding buffer: 20 mM Tris-Base pH 8.0, 250 mM NaCl, 5% glycerol and 20 mM imidazole; elution buffer: 20 mM Tris-Base pH 8.0, 250 mM NaCl, 5% glycerol and 500 mM imidazole). To remove the His-SUMO tag, the eluted proteins were concentrated in an Amicon®Ultra-15 Centrifugal Filter Unit (Millipore; 10 kDa cutoff) and incubated with the SUMO protease Ulp1 (Ulp1:His-SUMO-tH2 = 1:500 w/w) at 4 °C for 16 h. The tag-free tH2 was isolated by applying the sample to a HisTrap™ excel column to collect the flowthrough before being further purified with a Superdex-75 size-exclusion column in gel filtration buffer (20 mM Tris-Base pH 8.0, 250 mM NaCl, 5% glycerol).

### H2 protein structure prediction by AlphaFold2

We used the AlphaFold2 platform (https://colab.research.google.com/github/sokrypton/ColabFold/blob/main/AlphaFold2.ipynb) for H2 protein structure prediction [42]. The full-length H2 amino acid sequence was used as the query sequence, with the msa_mode of MMseqs2 (UniRef+Environmental) and a pair_mode of unpaired+paired. The predicted model with a high-confidence pIDDT score (>80) was considered as the candidate.

### Determination of the crystal structure of the H2 ectodomain (tH2, aa91-189)

For crystallization, 1 μl of tH2 (15 mg/ml) was mixed with 1 μl of the crystallization reagent from a reservoir containing 0.1 M sodium chloride, 0.1 M bicine pH 9.0, and 20% (w/v) PEG 550 MME. Crystals were obtained after two weeks of incubation at 20 °C. Ethylene glycol with a final concentration of 15% was used as a cryoprotectant for data collection. X-ray diffraction data of crystals were collected from beamline 15A, Taiwan Light Source (TLS), National Synchrotron Radiation Research Center in Hsinchu (NSRRC), Taiwan. The collected data were processed in HKL2000 [43] (program package of Denzo, XdisplayF, and Scalepack). The structure of tH2 was determined by molecule replacement in the program Molrep [44], with the AlphaFold2-predicted H2 structure used as the search model for molecule replacement. The program Refmac [45] was used to generate initial electron density maps by rigid-body refinement. For subsequent model building and refinement, the Refmac [45], Buccaneer [46], Coot [47], and PHENIX [48] programs were utilized. The data collection, refinement and structural statistics for tH2 are presented in Table 1. The UCSF Chimera program (https://www.cgl.ucsf.edu/chimera/) was used for structural analysis and figure generation [49, 50]. The structure factors and atomic coordinates of tH2 have been deposited in the Protein Data Bank (www.rcsb.org). The PDB ID of tH2 is 8INI.

**Table 1.** Data collection and refinement statistics for tH2 crystals.

|  |  |  |
| --- | --- | --- |
| Data collection | Native crystal | 488 |
| Wavelength (Å) | 1.00000 | 489 |
| Space group | *C2221* |  |
| Unit cell *a*, *b*, *c* (Å) | 46.7, 60.4, 131.6 | 490 |
| $\alpha$, $\beta$, $\gamma$ (°) | 90, 90, 90 | 491 |
| Resolution (Å) | 20-1.75 (1.81-1.75) | 492 |
| Unique reflections | 19019 (1895) | 493 |
| Redundancy | 3.6 (3.6) |  |
| Completeness (%) | 98.9 (100) | 494 |
| I / $\sigma$(I) | 34.2 (3.3) | 495 |
| R<sub>merge</sub> (%) | 5.6 (42.7) | 496 |
| Refinement |  |  |
| R<sub>work</sub> (%) | 16.9 | 497 |
| R<sub>free</sub> (%) | 20.3 | 498 |
| Bond r.m.s.d. (Å) | 0.007 | 499 |
| Angle r.m.s.d. (°) | 1.01 |  |
| Mean B value / no of atom | 25.8 / 1814 | 500 |
| Ramachandran plot (%) |  | 501 |
| Most favored (%) | 95.7 | 502 |
| Allowed (%) | 4.3 | 503 |
| Outliers (%) | 0 |  |
|  |  | 504 |

### Isothermal Titration Calorimetry (ITC) analyses of recombinant tH2 protein binding to truncated A28 protein *in vitro*

ITC analyses were performed using a MicroCal ITC200 isothermal titration calorimeter (Malvern Panalytical). All ITC assays were conducted using MES buffer (20 mm MES buffer pH 6.5 containing 50 mm NaCl). Recombinant A28 (aa56-146; sA28) protein was purified from *E. coli* as described previously [51], buffer-exchanged in the ITC buffer, and filtered through a 0.2-micron sterile filter. The tH2 and sA28 proteins were used at concentrations of 0.05 and 0.5 mM, respectively, for ITC measurements. During ITC assays, the degassed tH2 samples were maintained at room temperature (25 °C) and stirred at 300 rpm inside the ITC cell. For each titration, 2-μl aliquots of the sA28 protein were delivered into the tH2 sample at 150-s intervals to allow complete equilibration. The final ratio for all protein-protein interactions reached 1:1 at the end of the titration. Heat transfer was measured as a function of elapsed time. The corrected titration curve was fitted with a single-site binding model, and the thermodynamic parameters were calculated using the Origin software (version 7.0) provided by Malvern Panalytical Inc.

### Trans-complementation assays

Trans-complementation assays were performed as described previously [33, 40] with some modifications. In brief, freshly confluent 293T or BSC40 cells in a 6-well plate were transfected with 0.05 μg of wild-type or mutant H2R plasmids using 10 μl of lipofectamine 2000. At 24 h post-transfection, the cells were infected with vH2i at a multiplicity of infection (MOI) of 5 PFU per cell for 1 h at 37 °C and then incubated in complete growth media without IPTG. At 24 h post-infection (hpi), the whole cell lysates were collected, washed, and subjected to immunoblot analyses, plaque assays, and MV-induced membrane fusion assays as described below. Experiments with each H2 plasmid were repeated three times independently.

### Co-immunoprecipitation of vaccinia H2 and other EFC components

After being transfected with plasmid and infected with virus (tf/inf) as described above, 293T cells were harvested, washed, and lysed on ice in 500 μl of lysis buffer consisting of 0.5% NP-40, 20 mM Tris pH 8.0, 200 mM NaCl, and a protease inhibitor cocktail comprising 2 μg/ml aprotinin, 1 μg/ml leupeptin, 0.7 μg/ml pepstatin, and 1 mM phenylmethylsulfonyl fluoride. Clarified lysates were then incubated with A28 antisera-conjugated protein A beads (GE Healthcare) at 4 °C for 4 h with rotation at 15 rpm. The samples were subsequently denatured in SDS-containing sample buffer at 95 °C for 5 min, separated on SDS-PAGE gels and transferred onto nitrocellulose membranes for immunoblot analyses using anti-EFC component antibodies, including rabbit anti-A16, anti-G9, anti-L1, anti-G3, anti-L5, anti-F9, anti-A21, anti-H3 and anti-D8, all of which have been described previously ([35] and references therein). Anti-A28, anti-H2, anti-J5 and anti-A21 rabbit sera were newly prepared by immunizing NZW rabbits with soluble recombinant A28 (aa56-146), H2 (aa91-189), J5 (aa68-133) or A21 (aa39-117) proteins using an immunization protocol described previously [10, 11].

### MV-triggered cell fusion-from-without at acidic pH

Cell-cell fusion induced by vaccinia MV was performed as described previously with slight modification [34, 35]. Freshly confluent HeLa cells expressing GFP or RFP at a ratio of 1:1 in a 96-well plate were pretreated with 40 μg/ml cordycepin at 37 °C for 1 h prior to infection. Mutant H2 virus-containing cell lysates, harvested from the trans-complementation assays described above, were used to infect the GFP- and RFP-expressing HeLa cells at 37 °C for 1 h. Cells were subsequently washed with warm PBS and then subjected to pH 7.0 or 5.0 buffer treatment at 37 °C for 3 min, respectively, before being washed and then incubated in complete media containing 40 μg/ml cordycepin for another 2 h. These cells were photographed using a Zeiss Axiovert fluorescence microscope, and the percentage of cell fusion was analyzed using Fiji software and calculated as (the image area of GFP^+^RFP^+^ double-fluorescent cells divided by that of single-fluorescent cells) × 100, as described previously [34, 35].

### Statistical analysis

Statistical analyses were performed using Student’s *t*-tests in GraphPad Prism v.9.4.1. Data are expressed as mean ± SD. For comparisons with wild-type control, p values were adjusted by the “fdr” method using the “p.adjust” function in R v.4.2.2. Adjusted p values were considered statistically significant at *p<0.05; **p<0.01; ***p<0.001.

## Results

### The overall structure of the vaccinia H2 ectodomain

We determined the crystal structure of vaccinia tH2 (aa91-189) by X-ray diffraction at a resolution of 1.85 Å (Table 1). The orthorhombic tH2 crystals displayed a space group of C222_1_ and comprised two protein molecules in the asymmetric unit. A lattice packing analysis indicated that the tH2 crystal does not associate in specific ways with its neighbors. The two monomers are virtually identical in their three-dimensional structure, with an RMSD of 0.35 Å for 81 pairs of Cα atoms, excluding the C-terminal tails. The tH2 protein folds into a central five-stranded β-sheet and three cladding α-helices (Figure 1A&1B). The β-sheet is stabilized by disulfide bonds and hydrophobic cores on both sides (Figure 1C). On one side, the disulfide bond of C102-C148 anchors α-helix 1 onto β-sheet 1, while on the other side, the disulfide bond of C162-C182 tethers helix α3 to the β5-α2 loop. Around these two disulfide bonds, many nonpolar amino acids (such as V103, F147, I160, W181, etc. in Figure 1C) constitute a stable hydrophobic core, rendering the overall structure more stable.

**Figure 1.**
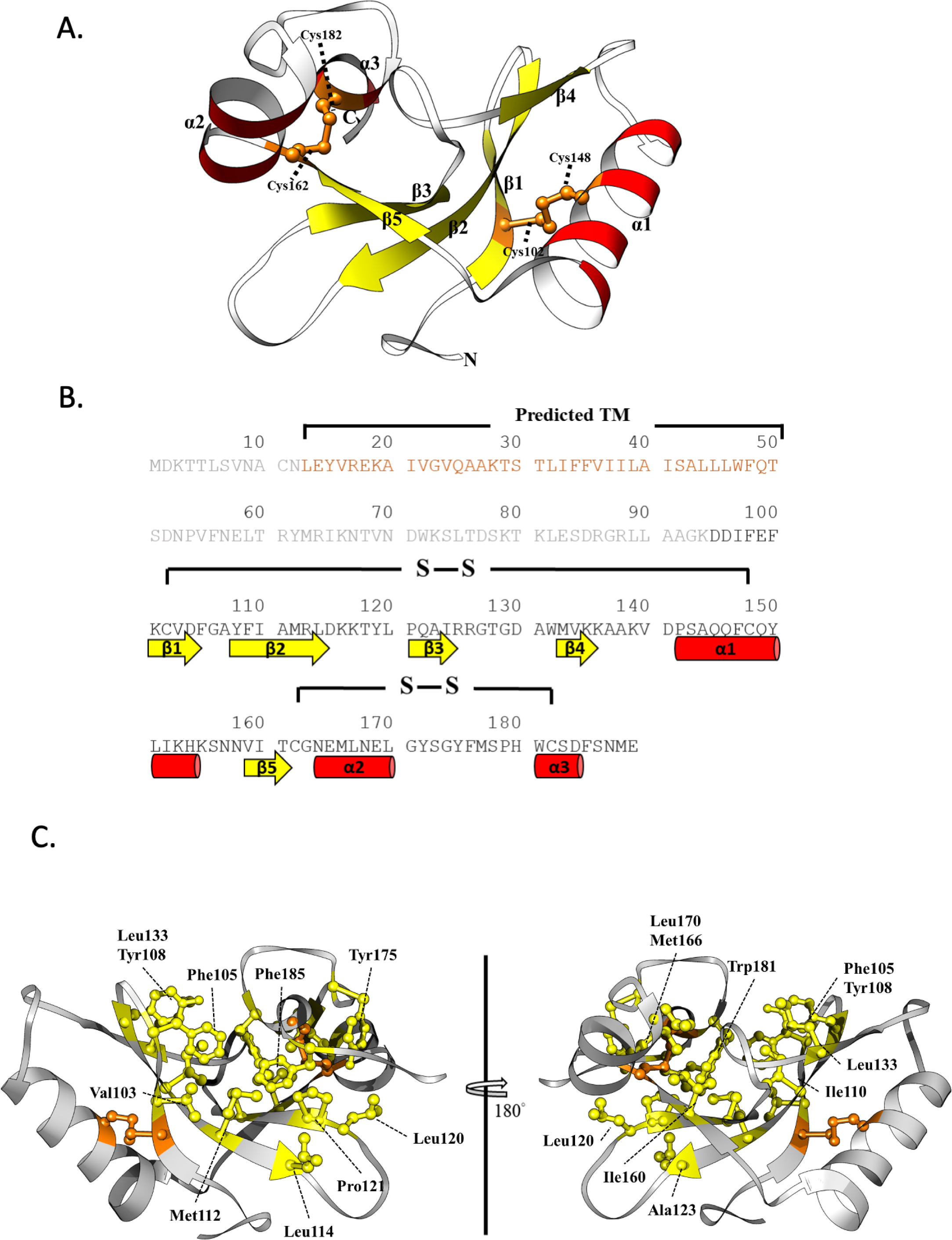
Crystal structure and sequence analysis of H2 protein. **(A)** Schematic diagram of the vaccinia virus tH2 structure (H2 ectodomain; aa95-189). A 3D flat ribbon model is presented, with loops, α-helixes and β-sheets in white, red and yellow, respectively. tH2ΔTM comprises three α-helices (α1, α2, α3) and five β-sheets (β1, β2, β3, β4, β5). Disulfide bonds between Cys (C)102 and C148, and between C162 and C182, are shown as orange sticks. **(B)** The amino acid sequence of the full-length H2 protein with residues colored-labeled as in (A). Residues in black font denote the tH2 ectodomain (aa95-189) that was subjected to crystallization experiments, whereas the residues in brown font (aa13-50) were predicted as a transmembrane (TM) domain by AlphaFold2. The residues in gray reflect the H2 flexible structure indicated by AlphaFold2. **(C)** The hydrophobic core of tH2. All the hydrophobic residues that contribute to the hydrophobic core are colored in yellow using the stick-ball model.

### Mutagenesis screen for residues of the tH2 ectodomain critical for viral infectivity, A28 interaction and EFC formation

We aimed to study the functional role of H2 in EFC-triggered membrane fusion using mutational analysis based on structural and genetic information. Resolving the atomic structure of the tH2 ectodomain allowed us to define surface-charged clusters potentially responsible for protein-protein interactions. We also targeted highly conserved residues based on a multiple sequence alignment of 28 poxviruses isolated from various animal species (Figure 2). To identify residues critical to viral growth, we assayed viral infectivity of all the H2 mutant constructs using a trans-complementation assay in 293T cells as described in the materials and methods. In brief, we transfected plasmids encoding either wild type (WT) or mutant H2 protein 24 h prior to infection with vH2i at an MOI of 5 without IPTG. Then, we harvested the transfection/infection (tf/in) lysate at 24 hpi and titered the viral yield on BSC40 cells in the presence of 50 μM IPTG. Considering that Nelson *et al*. [40] mapped the peptide sequence “LGYSG” as critical for viral infectivity, we generated two loss-of-function mutants (G^171^G^174^A and Y^172^A) as inactive mutant controls in our analysis. Overall, we designed an additional 26 plasmids harboring mutation(s) in the H2 ectodomain and all the mutant H2 proteins were successfully expressed (Figure 3A).

**Figure 2.**
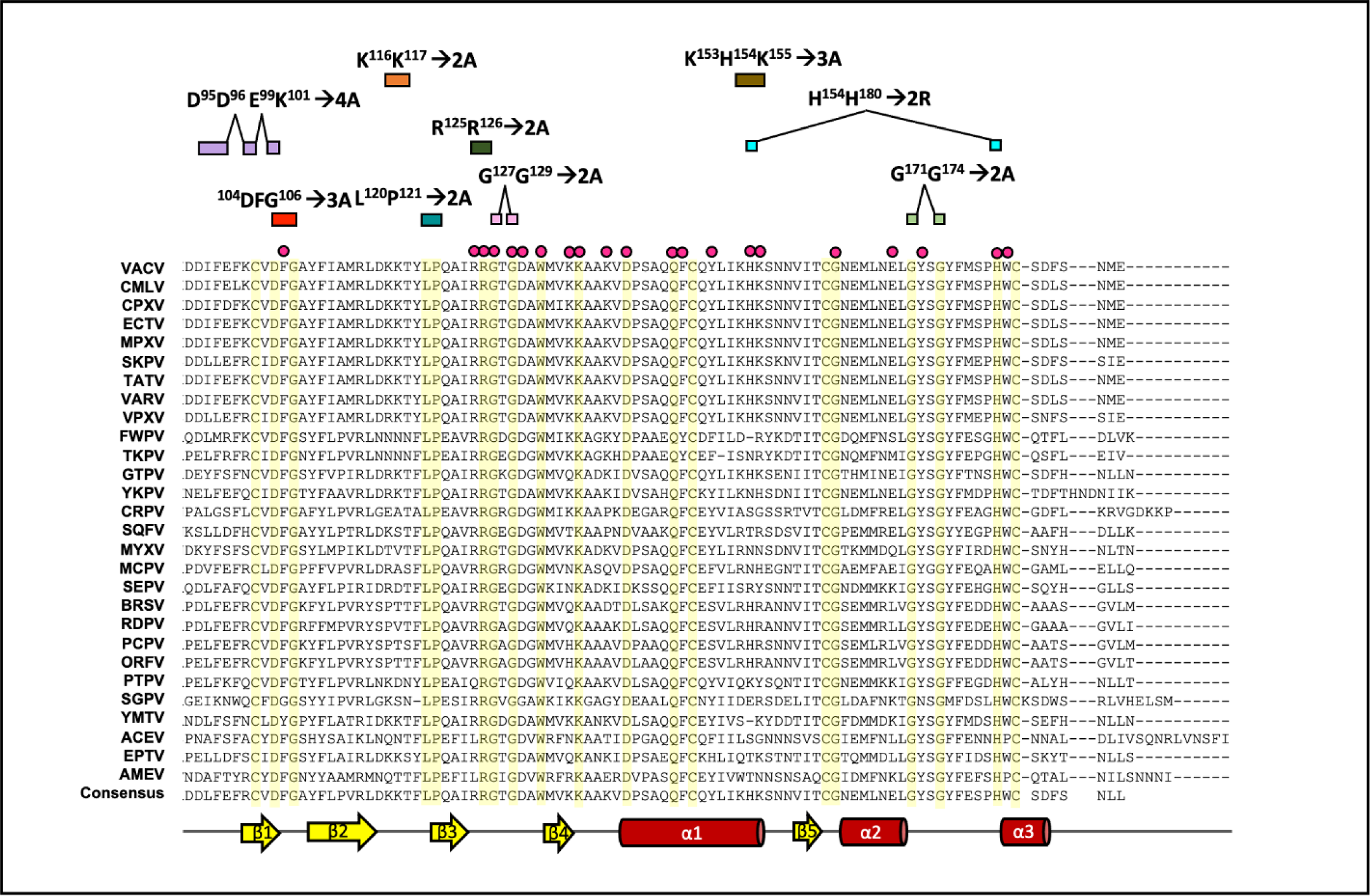
Multiple sequence alignment of the poxviral H2 ectodomain and mutant designations. The amino acid sequence of the crystallized vaccinia virus H2 C-terminal ectodomain (aa95–189) aligned with orthologs from 24 representative chordopoxviruses and three entomopoxviruses. Residues displaying 100% identity across the selected poxviruses are highlighted in yellow. The resolved secondary structures are shown at the bottom. Alanine mutations generated in this study are denoted as red circles (single mutation) or bridged boxes (clustered mutations) and are displayed above the alignment. VACV: vaccinia virus (YP_233033.1); CMLV: camelpox virus (NP_570537.1); CPXV: cowpox virus (NP_619947.1); ECTV: ectromelia virus (NP_671649.1); MPXV: monkeypox virus (NP_536567.1); SKPV: skunkpox virus (YP_009282841.1); TATV: taterapox virus (YP_717459.1); VARV: variola virus (NP_042179.1); VPXV: volepox virus (YP_009282895.1); FWPV: fowlpox virus (NP_039155.1); TKPV: turkeypox virus (YP_009177163.1), GTPV: goatpox virus (YP_001293309.1), YKPV: yokapox virus (YP_004821485.1); CRPV: crocodilepox virus (QGT49410.1); SQPV: squirrelpox virus (YP_008658542.1); MYXV: myxoma virus (NP_051830.1); MCPV: molluscum contagiosum virus (NP_044085.1); SEPV: sea otter poxvirus (YP_009480656.1); BPSV: bovine papular stomatitis virus (NP_958014.1); RDPV: red deer parapoxvirus (YP_009112844.1); PCPV: pesudocowpox virus (YP_003457411.1); ORFV: orf virus (NP_957882.1); PTPV: pteropox virus (YP_009268838.1); SGPV: salmon gill poxvirus (AKR04251.1); YMTV: yaba monkey tumor virus (NP_938373.1); ACEV: anomala cuprea entomopoxvirus (YP_009001544.1); EPTV: eptesipox virus (YP_009408076.1); AMEV: amsacta moorei entomopoxvirus (NP_064968.2).

**Figure 3.**
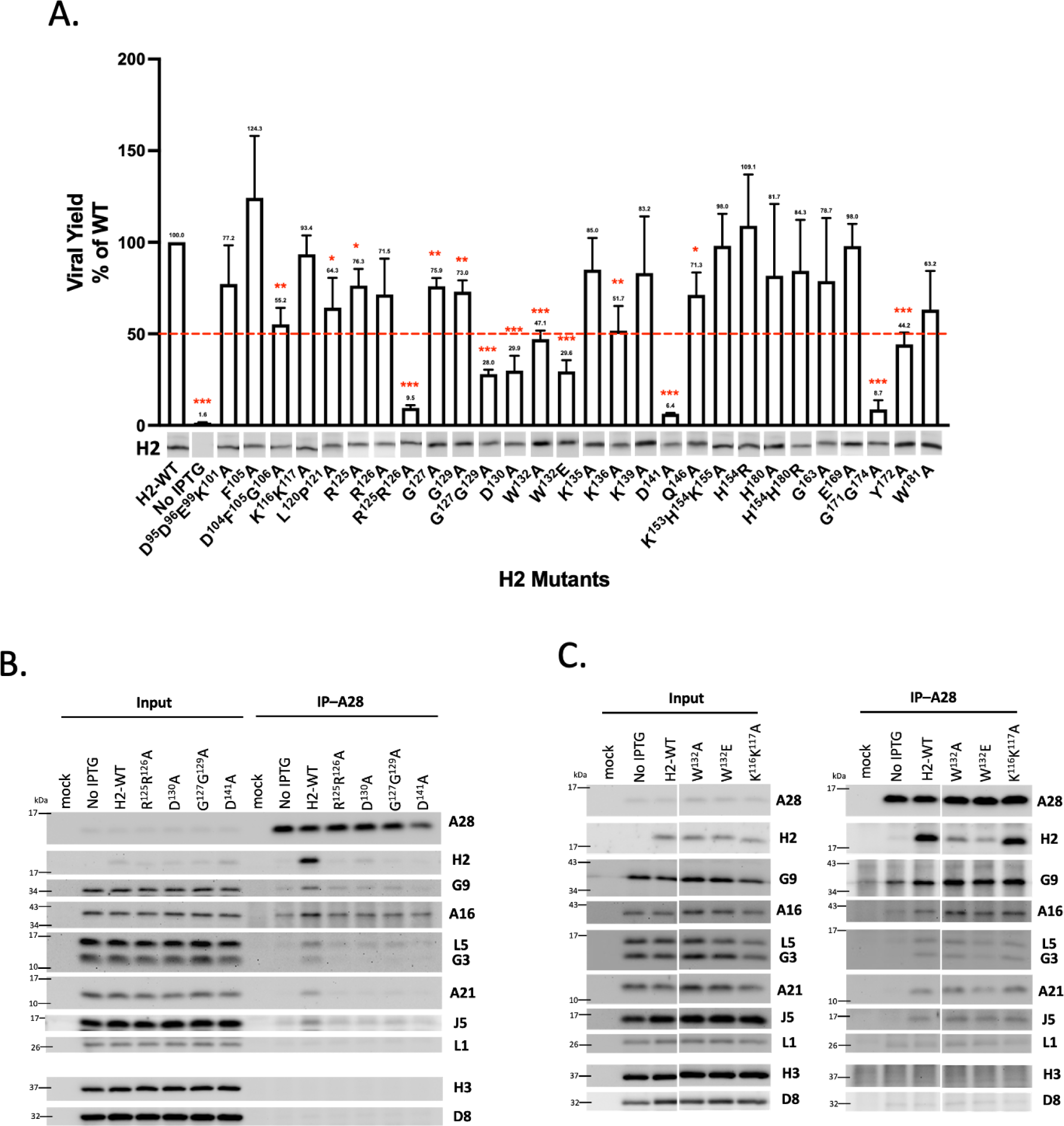
Biological effects of mutations in the ectodomain of vaccinia H2. **(A)** Trans-complementation assays of H2 mutants hosting mutations in the crystallized C-terminal ectodomain. HEK293T cells were transfected with vector, wild-type (WT) or mutant H2 plasmids and then infected with vH2i, before being harvested at 24 h post-infection. Plaque assays were performed to obtain the mutant virus titers, which were then normalized to that of the WT by dividing the titer of each mutant with that of the WT-H2. All mutants were assayed independently three times and the mean values are presented with standard deviation. The red dashed line represents 50%. H2 expression in crude lysate was detected by immunoblotting using an anti-H2 antibody. Student’s *t*-test: \**P* < 0.05, \*\**P* < 0.01, \*\*\**P* < 0.001. **(B & C)** Co-immunoprecipitation (co-IP) of A28 protein with various H2 mutants. Trans-complementation lysates were incubated with anti-A28-conjugated agarose beads, washed, and then the immunoprecipitates were analyzed by immunoblotting using antibodies against anti-EFC components. Anti-H3 and anti-D8 antibodies served as negative controls.

Viral yields of the two loss-of-function H2 mutants (G^171^G^174^A and Y^172^A) were significantly reduced, consistent with the previous report [40]. For the other 26 H2 mutant constructs, six mutants—namely R^125^R^126^A, G^127^G^129^A, D^130^A, W^132^A, W^132^E and D^141^A—produced viral yields less than 50% of the WT (Figure 3A). Notably, we titrated the mutant H2 plasmids to obtain a comparable protein expression level in our tf/if experiments; therefore, the reduced virus titers for the mutants were not due to different levels of H2 protein expression.

H2 is known to interact with A28 in the absence of other EFC components [21]. Accordingly, we selected our H2 mutants, R^125^R^126^A, D^130^A, G^127^G^129^A, D^141^A, W^132^A and W^132^E, for which virus titers were reduced by more than 50% for co-immunoprecipitation (co-IP) using the cell lysates harvested from the trans-complementation assays (Figure 3B & 3C). We also included WT-H2 (Figure 3B) and an active H2 mutant, K^116^K^117^A, as a co-IP control (Figure 3C). In the co-IP analyses, we used rabbit anti-A28 antibody to immunoprecipitate A28 protein and analyzed the amounts of A28-bound H2 protein and other A28-bound EFC components in immunoblot analyses. As shown in Figure 3B, four of the H2 mutants—R^125^R^126^A, D^130^A, G^127^G^129^A, and D^141^A—could not bind A28 protein. Moreover, the other EFC components either were barely detected in the co-IP or displayed a reduced signal intensity, i.e., comparable to the background level in the negative control (no IPTG), suggesting that these mutations not only interrupted A28-H2 interaction but also affected assembly of the whole EFC. As shown in Figure 3C, two H2 mutants, W^132^A and W^132^E, also bound less well to A28 relative to both WT and the K^116^K^117^A active mutant. Interestingly, levels of other EFC components brought down by A28 antibody remained largely unaltered for these two mutants (Figure 3C), suggesting that mutations of W132 had a more local effect in A28-H2 interaction with little influence on EFC assembly.

To verify that these H2 mutations directly affect H2 interaction with A28, we performed ITC assays using purified recombinant wild type sA28 (aa56-146) and the various tH2 (aa91-189) mutant proteins. As shown in Figure 4A&B, the active tH2 mutant K^116^K^117^A bound to sA28 protein with a dissociation constant K_D_ ∼ 5.3 μM, i.e., similar to our recent finding for wild type tH2 protein [51], whereas all five defective H2 mutants, R^125^R^126^A, D^130^A, G^127^G^129^A, D^141^A and W^132^A, displayed significantly reduced binding ability to sA28, with K_D_ values of ∼0.18-8.9 mM. Taken together, these results reveal that the R125, R126, G127, G129, D130, D141 and W132 residues are important for binding of proteins H2 to A28, both *in vivo* and *in vitro*.

**Figure 4.**
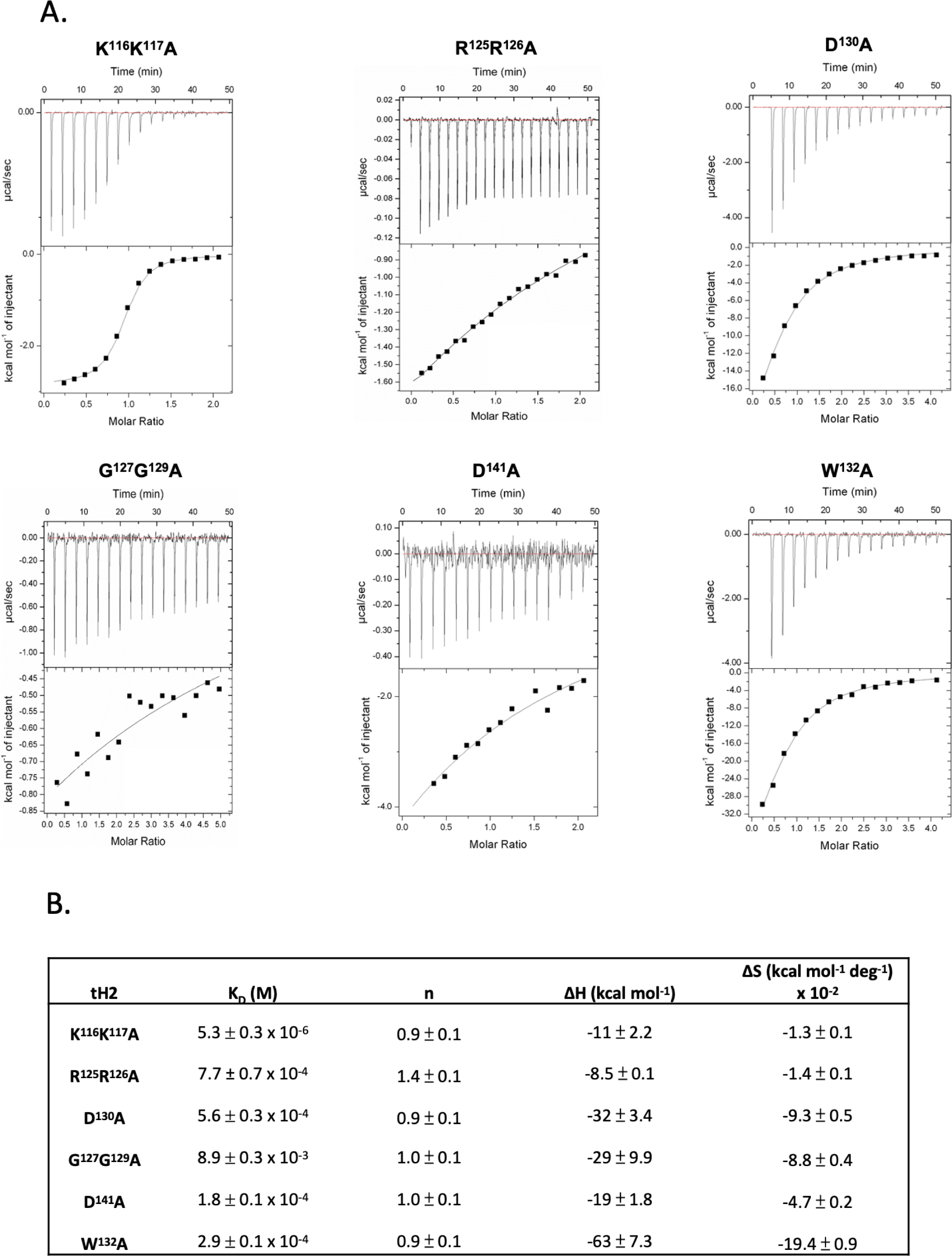
*In vitro* ITC analyses of mutant tH2 protein binding to wild-type sA28 protein. **(A)** Recombinant vaccinia sA28 (0.5 mM) was isothermally titrated with various vaccinia tH2 mutant proteins (0.05 mM), including R^125^R^126^A, D^130^A, G^127^G^129^A, D^141^A and W^132^A. The active tH2 mutant protein K^116^K^117^A was included as a positive control. The ITC exothermic heat of the reaction (upper panel) and the integrated areas of the peaks (lower panel) plotted against the molar ratio of the two proteins are shown. The best fit of the experimental data was determined from the non-linear least square fit (solid line). **(B)** Thermodynamic parameters derived from the ITC data of protein-protein interactions shown in (A).

Based on our resolved tH2 structure (Figure 1), the R125, R126, G127, G129, D130 and W132 residues reside in loop 125-132 flanked by β3 and β4. R125 forms a salt bridge with E169 and generates additional H-bonds with T128 and G129 (Figure 5A), which provide structural stability between loop 125-132 and α2. The protonated guanidinium group of R126 contacts the indole side chain of W132 using cation–π interaction [52]. Together, these interactions shape the conformation of loop 125-132 (Figure 5A). Residue D130 makes H-bonds with Y172 and W181 to support the association of loop 125-132 with α1 and α2 (Figure 5B). Accordingly, mutating R125, R126 and D130 to alanine disrupts these interactions, destabilizing the conformation of loop 125-132. Both G127 and G129 are fully conserved across the poxvirus family (Figure 2). Since glycine residues in turns or short loops often contribute to chain compaction during the early folding state [53], substituting those glycine residues to alanine may disfavor the proper bending of loop 125-132 (Figure 5C). Interestingly, the location of W132 is at the edge of loop 125-132, not the center. The indole ring of W132 is incorporated into other hydrophobic side chains of β4 and α1 residues (V134, F147, and L151) to form a hydrophobic core [54, 55] that, in turn, confines the organization of loop 125-132, and the β4 and α1 regions (Figure 5D). In addition, W132 exhibits an unusual solvent-exposed propensity and, when combined with our co-IP and ITC data, we infer that W132 uses its bulky hydrophobic indole ring to interact directly with the H2 binding partner A28. The W^132^E mutant displayed a worse complementation phenotype than observed for W132A (Figure 3A), indicating that the negatively charged carboxylate group further disrupts the local structure to aggravate H2 function. Taken together, our results revealed that loop 125-132 is an important domain for viral infectivity, H2-A28 interaction and subsequent EFC assembly.

**Figure 5.**
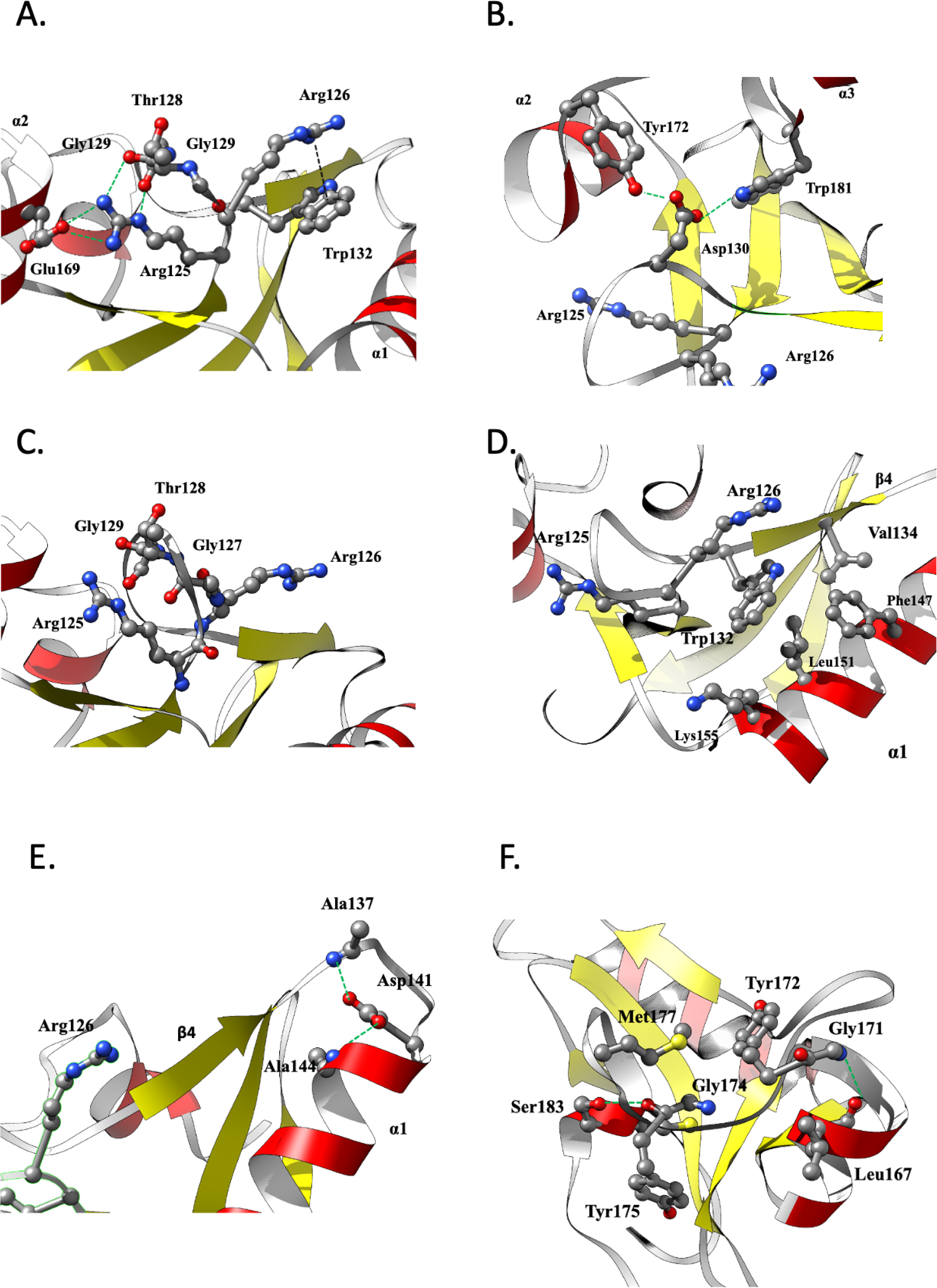
Structural insights in residues important for vaccinia MV infectivity based on mutagenesis assay. The tH2 protein structure, as presented in the flat ribbon cartoon model, with relevant residues shown in ball and stick format. Dashed lines label interactions between two molecules. **(A)** The loop 125-132 region with Arg (R)125/126, for which double mutation reduced vaccinia virus infectivity by 90%. Residue R125 interacts with Thr (T)128, Gly (G)129 and Glu (E)169 (H-bonds are colored green), whereas residue R126 interacts with Trp (W)132 (cation–π interaction colored black). **(B)** The loop 125-132 region with Asp (D)130, for which a single mutation reduced infectivity by 70%. Residue D130 interacts with Tyr (Y)172 and W181 (H-bonds are colored green). **(C)** Positions of residues G127 and G129 on loop 125-132 with respect to surrounding residues. Double mutation of G127/129 reduced virus infectivity by 70%. **(D)** The loop 136-141 region with residue D141 and its interacting partners Ala (A)137 and A142. Single mutation of D141 reduced virus infectivity by 90%. **(E)** The hydrophobic core region comprised of W132, Val (V)134, Phr (F)147, Leu (L)151 and Lys (K)155. Single mutation of W132 reduced virus infectivity by 70%. **(F)** The loop 171-180 region with G171 and G174 and surrounding residues L167, Y172, Y175, Met (M)177 and Ser (S)183. The H-bond between G171 and L167 is colored green. Double mutation of G171/174 reduced virus infectivity by 90%.

Besides mutations in the loop 125-132, the D^141^A mutation also affected H2 function. Negatively charged residue D141 on the opposite side of β4 makes H-bond interactions with the main-chain nitrogens of A137 and A144 (Figure 5E). These interactions facilitate the formation of loop 136-141, provide structural stability for the β4 and α1 association, and indirectly shape the conformation of loop 125-132. Therefore, as observed for the residues constituting loop 125-132, alanine mutation of D141 deforms the critical loop 125-132, thereby negatively affecting H2 function and viral infectivity.

The G^171^G^174^A double mutant displayed significantly reduced MV infectivity (Figure 3A), as reported previously [40]. Similar to G127 and G129 (Figure 5C), both G171 and G174 are located in a loop between α2 and 3 (Figure 5F). Thus, the glycine residues in the loop regions of H2 may play an important role in the protein’s structural folding, stability, and function.

### Hydrophobic residues and charged clusters in the H2 N-terminal are crucial for vaccinia virus infectivity and EFC formation

The tH2 structure we resolved does not include the N-terminal region (aa1-90). To investigate the structural details of this region, we modeled full-length H2 using the AlphaFold2 (AF2) [42] online package and then fitted our crystal structure within the predicted H2 protein structure. We generated a high-confidence AF2-predicted H2 structure, as supported by the high pIDDT score of >80. Superimposition of the crystal structure and *in silico*-generated model revealed an RMSD value of 0.5 Å, further supporting the high accuracy of the predicted full-length H2 structure.

The AF2-predicted structure revealed a long α-helix (aa55-90) that linked the putative N-terminal transmembrane helix (aa13-50) to the tH2 ectodomain (aa91-189) via short loops at both ends. To investigate the role of the hydrophobic N-terminal region (shown as a light blue bar at the bottom of Figure 6A), we conducted alanine mutagenesis on the fully conserved S42 and I65 residues, as well as on residues within the long α-helix (E58, R64, K66, W72 and K79) and in the E^83^S^84^D^85^R^86^G^87^R^88^ charged cluster (Figure 6A). Trans-complementation assays on these H2 mutant constructs revealed that the I^65^A single mutant displayed a >50% reduction in viral infectivity. When combined with mutations of neighboring conserved residues R64 and K66 (mutant R^64^I^65^K^66^A) or of hydrophobic W72 (mutant I^65^W^72^A) that extends its indole ring parallel to the I65 side-chain, viral infectivity dropped to less than 10%. The virus titer of the E^83^S^84^D^85^R^86^G^87^R^88^A mutant was also less than 50% of the WT titer (Figure 6B). Remarkably, co-IP using anti-A28 antibody revealed that A28 protein remained associated with the I^65^A, R^64^I^65^K^66^A, I^65^W^72^A and E^83^S^84^D^85^R^86^G^87^R^88^A mutant H2 protein but not with other EFC components (Figure 6C). Although G9 and A16 proteins were also detected in these mutant co-IP samples, both band intensity was comparable to that of the negative control sample (i.e., no IPTG). Taken together, these data reveal that mutations in the N-terminal α-helix (aa55-90) of the H2 protein interfere with EFC formation and virus infectivity without affecting its binding to A28.

**Figure 6.**
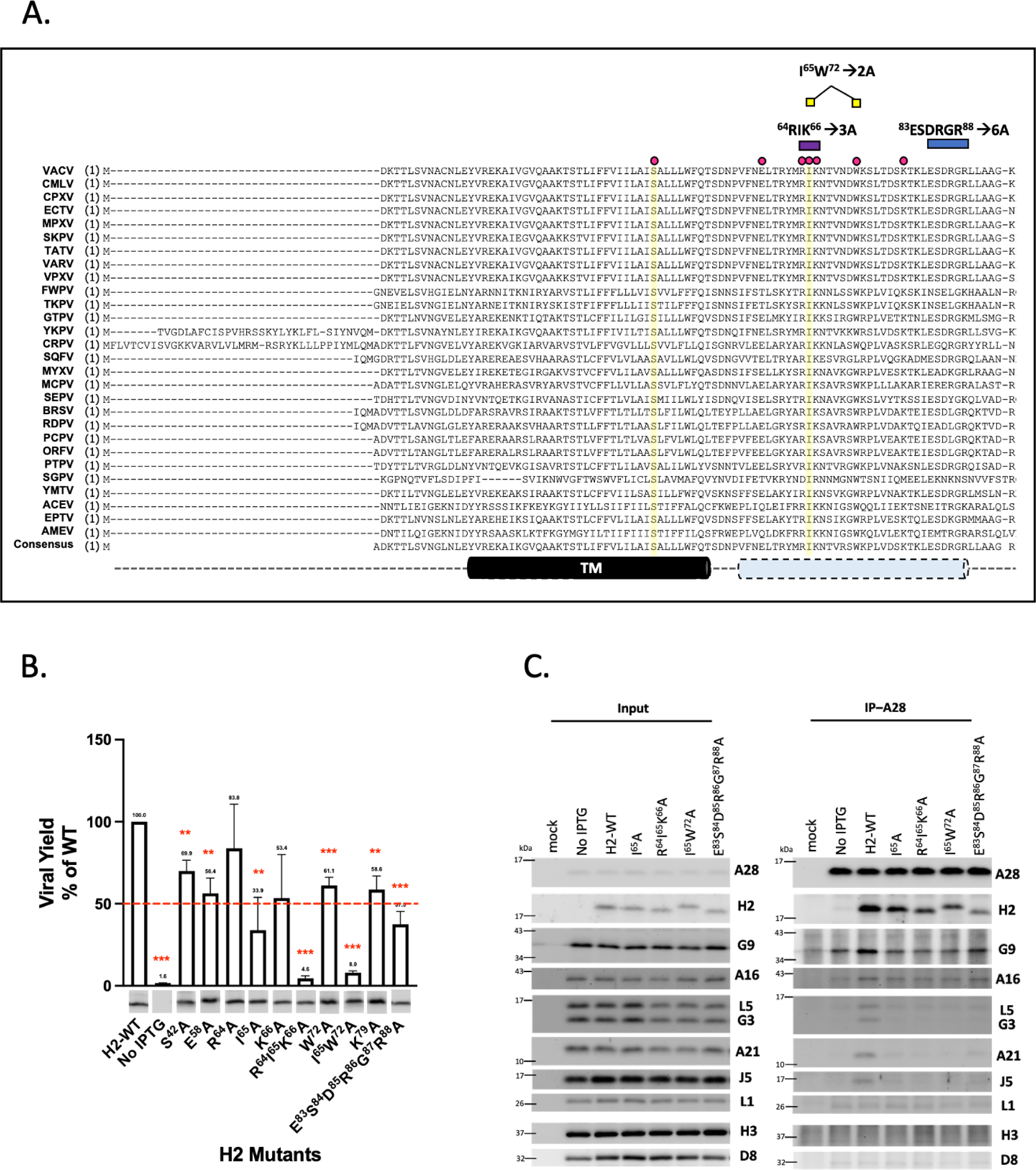
Mutations of the H2 N-terminal α-helical region and their biological effects. **(A)** Alignment of orthologous poxviral H2 sequences at the N-terminus and mutant designations. Residues displaying 100% identity across the selected poxviruses are highlighted in yellow. The predicted α-helical structures (aa55-90, light blue) and the transmembrane (TM) region (aa13-50, black) are shown at the bottom of the figure. Alanine mutations generated in this study are denoted as red circles (single mutation) or bridged colored boxes (clustered mutations) and are displayed above the alignment. VACV: vaccinia virus (YP_233033.1); CMLV: camelpox virus (NP_570537.1); CPXV: cowpox virus (NP_619947.1); ECTV: ectromelia virus (NP_671649.1); MPXV: monkeypox virus (NP_536567.1); SKPV: skunkpox virus (YP_009282841.1); TATV: taterapox virus (YP_717459.1); VARV: variola virus (NP_042179.1); VPXV: volepox virus (YP_009282895.1); FWPV: fowlpox virus (NP_039155.1); TKPV: turkeypox virus (YP_009177163.1), GTPV: goatpox virus (YP_001293309.1), YKPV: yokapox virus (YP_004821485.1); CRPV: crocodilepox virus (QGT49410.1); SQPV: squirrelpox virus (YP_008658542.1); MYXV: myxoma virus (NP_051830.1); MCPV: molluscum contagiosum virus (NP_044085.1); SEPV: sea otter poxvirus (YP_009480656.1); BPSV: bovine papular stomatitis virus (NP_958014.1); RDPV: red deer parapoxvirus (YP_009112844.1); PCPV: pesudocowpox virus (YP_003457411.1); ORFV: orf virus (NP_957882.1); PTPV: pteropox virus (YP_009268838.1); SGPV: salmon gill poxvirus (AKR04251.1); YMTV: yaba monkey tumor virus (NP_938373.1); ACEV: anomala cuprea entomopoxvirus (YP_009001544.1); EPTV: eptesipox virus (YP_009408076.1); AMEV: amsacta moorei entomopoxvirus (NP_064968.2). **(B)** Complementation assays of H2 mutants involving mutations in the N-terminal α-helical region. HEK293T cells were transfected with plasmids expressing either vector, wild-type (WT) or mutated H2 and then infected with vH2i, before being harvested at 24 h post-infection. The virus titers were determined by plaque assays and normalized by dividing the titer for each H2 mutant with that of the WT. All assays were repeated three times and mean values are shown with standard deviation. Student’s *t*-test: \**P* < 0.05, \*\**P* < 0.01, \*\*\**P* < 0.001. The red dashed line marks the 50% viral yield of the WT. H2 expression in crude lysate was detected by immunoblotting using an anti-H2 antibody. (C) Co-immunoprecipitation (co-IP) of defective α-helical H2 mutants. Cell lysates from transient complementation assays were prepared and then incubated with anti-A28-conjugated agarose beads before analyzing the immunoprecipitates using individual anti-EFC antibodies. Anti-H3 and anti-D8 antibodies served as negative controls.

### Loss-of-function H2 mutants exhibit defective membrane fusion activity

A previous fluorescence-dequenching experiment showed that H2-deficient vaccinia MV particles failed to trigger hemi-fusion with the host cell membrane [28]. To explore if the critical residues we identified in our trans-complementation assays described above also contribute to fusion activity, we performed mature virion (MV)-triggered cell fusion assays on lysates harvested from the trans-complementation assays. Vaccinia morphogenesis is not affected by the deletion of EFC components ([27] and references therein). In a parallel study of vaccinia virus A28 protein function, we also confirmed that mutations of vaccinia A28 specifically affect MV particle infectivity without impairing MV morphogenesis [51], supporting the feasibility of conducting MV-triggered cell fusion assays using the lysates generated from our trans-complementation assays.

HeLa cells expressing GFP or RFP were mixed at a 1:1 ratio and infected with lysates containing WT or mutant H2 viruses, treated with neutral pH 7 or acidic pH 5 buffer, and MV-triggered cell-cell fusion was monitored, as described in materials and methods. As expected, no cell-cell fusion was triggered by WT or mutant H2 viruses at neutral pH (Figure 7A, top panel). Whereas WT and active mutant K^116^K^117^A triggered robust cell-cell fusion under acidic conditions as expected, all of the H2 mutant lines that had displayed reduced viral growth and impaired binding ability to A28 protein and/or other EFC components also exhibited reduced MV-induced membrane fusion activity at low pH (Figure 7A, bottom panel). Relative quantification of the above MV-triggered cell fusion is shown in Figure 7B and revealed a consistent pattern between viral yield and MV-triggered membrane fusion (Figure 7C) for all H2 mutants we examined. Thus, we have identified specific residues in the N-terminal region and the ectodomain of vaccinia H2 protein that are important for EFC-mediated membrane fusion function.

**Figure 7.**
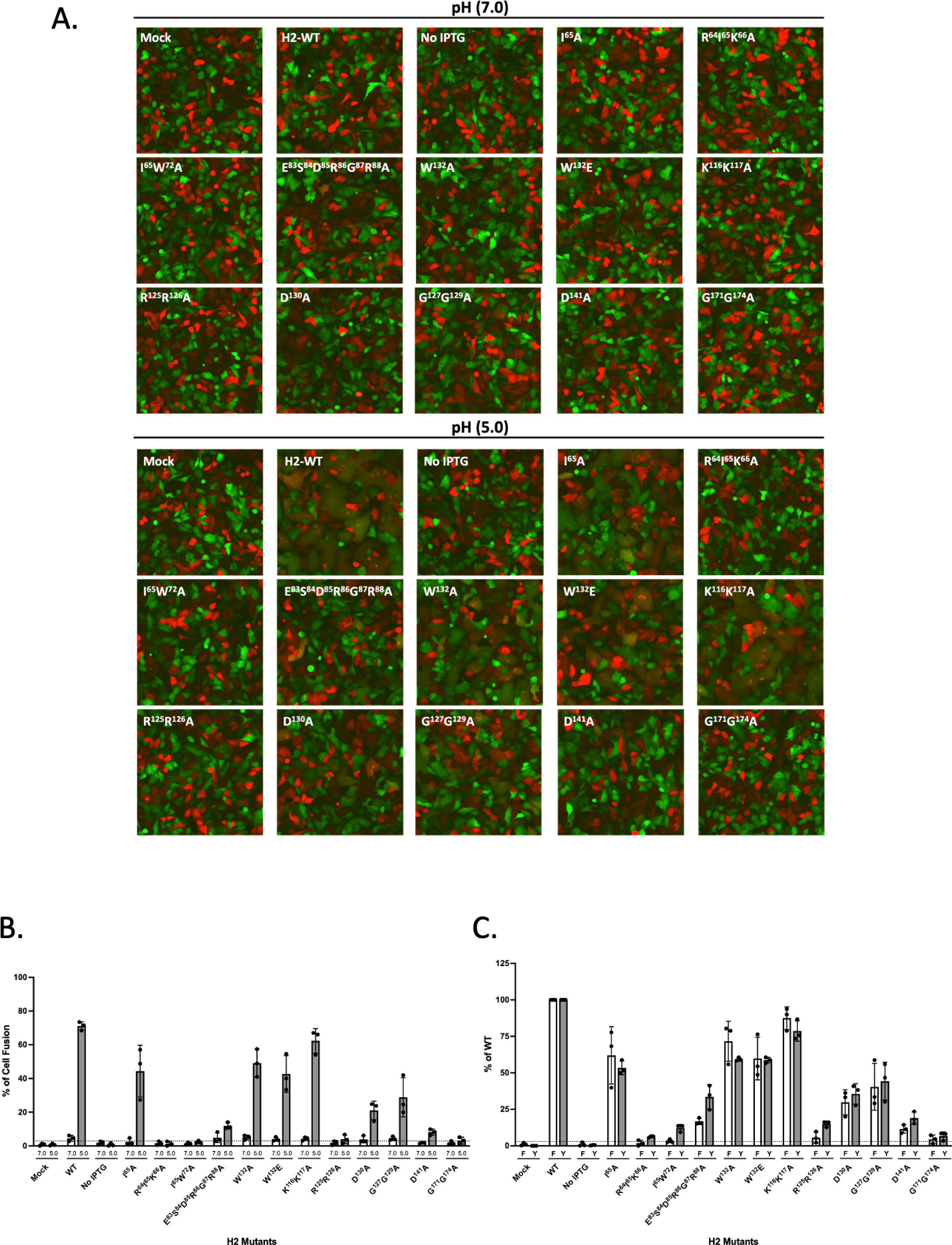
Cell-cell fusion assays of H2 mutants reveal impaired fusion at low pH. **(A)** HeLa cells expressing GFP or RFP were mixed at a ratio of 1:1 and treated with lysates containing WT or mutant H2 viruses in neutral pH7 or acidic pH5 buffers. Fluorescent cell images were photographed at 2 hpi. **(B)** Quantification of MV-induced cell-cell fusion at pH7 and pH5 from the images in (A). Photographs from three independent experiments were analyzed using Fiji software and the percentages of virus-triggered cell fusion at pH 5 were calculated as (the surface area of GFP^+^RFP^+^ double-fluorescent cells divided by the surface area of single-fluorescent cells) x100%. Error bars denote standard deviations from three independent experiments. The dashed line represents the averaged background fusion signal induced by H2 mutants at pH 7. Statistical comparisons of cell-cell fusion are between WT and each mutant at the low pH of 5. Student’s *t*-test: \**P* < 0.05, \*\**P* < 0.01, \*\*\**P* < 0.001. ns: not significant. **(C)** Comparison of MV-triggered cell fusion at low pH (F) and 24h viral yield (Y) after normalized to those of WT. H2 mutants with impaired viral infectivity (<50% of WT) displayed comparable fusion defects.

Finally, to provide a 3D illustration of these critical residues that we identified and characterized in this study, we have colored them in our crystal structure of the H2 ectodomain (Figure 8, top), as well as in the full-length H2 AlphaFold-generated model structure (Figure 8, bottom), in yellow, orange and red to reflect structural regions of increased functional importance (as described in the legend of Figure 8). Combining the results of our structural and functional analyses, we assert that the surface of the ectodomain, including two loop regions, ^170^LGYSG^174^ and ^125^RRGTGDAW^132^, constitutes a broad A28-binding region. Moreover, although not involved in A28 binding, the N-terminal helical region approximal to the membrane, encompassing ^64^RIK^66^, ^72^W and ^83^ESDRGR^88^, is also important for viral EFC formation and MV infectivity.

**Figure 8.**
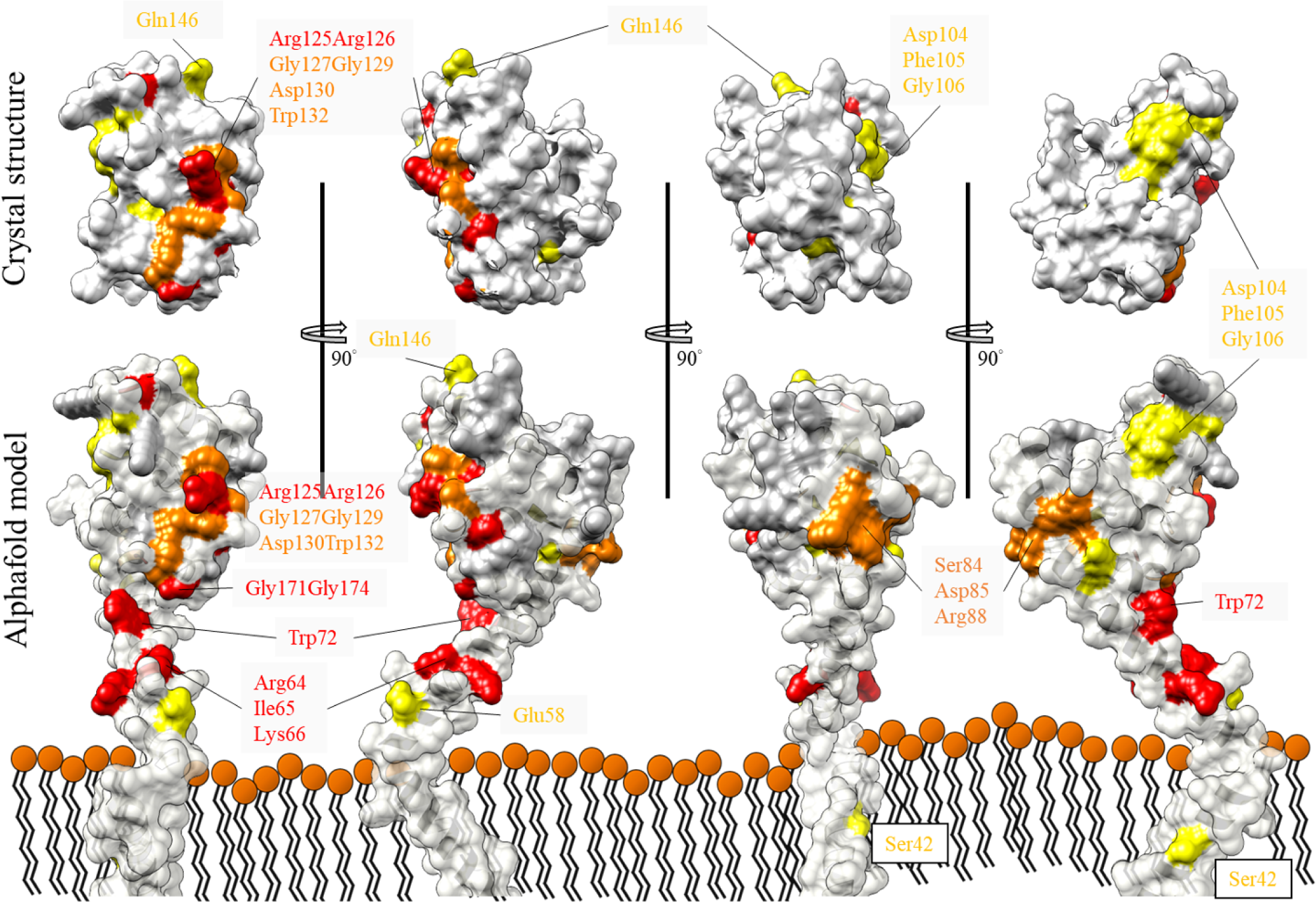
Summary of important residues for H2 protein on the 3D modeled structure. Highlight functional important residues in the tH2 crystal structure (top row) and the Alphafold2 predicted full-length H2 model (bottom row), shown in the 3D surface structure. A 360° lateral view of the tH2 crystal structure and the full-length H2 AlphaFold2-predicted model colored with residues that, when mutated, exhibited a significant loss of MV infectivity: mutant titer < 20% of WT (in red), ∼20-50% of WT (in orange), ∼50-75% of WT (in yellow). Ninety-degree view of the 3D surface structure rotated clockwise.

## Discussion

Enveloped viruses encode viral fusion proteins to trigger virus-cell membrane fusion and initiate the infectious cycle. In recent decades, the resolution of various viral fusion protein structures has contributed invaluable knowledge toward our understanding of the molecular process of membrane fusion induced by conformational changes of viral fusion proteins [1, 2, 56–58]. Despite having distinct structural features that classify known viral fusion proteins into three groups, the various viral fusion proteins display mechanistic conservation [59]. First, all viral fusion proteins undergo a cascade of structural rearrangements to bridge and ultimately merge the opposing viral and host membranes. Second, all viral fusion proteins carry a shielded hydrophobic fusion peptide or fusion loop for insertion into the host lipid bilayer upon acid-induced conformational changes. Third, some viral fusion proteins contain a membrane-proximal segment that is rich in hydrophobic and polar residues, termed the membrane-proximal (external) region, which regulates fusion activity by interacting with viral envelope lipids and/or adjacent protomers or by shielding the fusion loop [60, 61]. Notably, a recent report of a prokaryotic virus expressing a fusion protein with a novel fold [62] has raised the intriguing questions of which vaccinia EFC components harbor a fusogenic structural element and how these 11 components undergo extensive conformational changes to execute membrane fusion.

H2-deficient vaccinia MV cannot proceed to hemifusion upon membrane attachment [28], suggesting that it participates in the first step of the fusion process. Nelson *et al*. [40] identified a conserved fusion peptide-like sequence, ^170^LGYSG^174^, flanked by cysteine residues in the H2 C-terminal domain important for binding to A28 and MV infectivity. The current study further identified the ^125^RRGTGDAW^132^ sequences as critical to H2-A28 interaction, virus infectivity and membrane fusion activity. Interestingly, both sequences harbor two fully conserved glycines and a hydrophobic tryptophan. Based on our tH2 structure, both ^170^LGYSG^174^ and ^125^RRGTGDAW^132^ form flexible loops that project toward the same side of the protein, together contributing to a surface for binding H2 to A28 (Figure 8). This scenario is largely consistent with a molecular dynamics simulation of soluble A28 (aa56-146) and H2 (aa91-189) from a parallel study of vaccinia A28 protein [51], which revealed that these two loops make extensive contact with A28 via both electrostatic and polar interactions. Thus, based on our analyses and those of the Nelson *et al*. study [40], each loop is necessary but insufficient for the H2-A28 interaction, with loop disruption eliciting defects in virus-host membrane fusion.

Between the crystallized tH2 ectodomain (aa91-189) and the putative transmembrane region (aa13-50), computational modeling predicted a long α-helix (aa51-90) connecting both ends with short loops. Based on a recent lipid-embedded H2-A28 subcomplex computer simulation [51], we proposed that this distinct structural domain lies on the top of the viral envelope and appears intimately associated with the outer leaflet. In the current study, the variants hosting mutations in this region that we observed as eliciting impaired viral growth, i.e., I^65^A, R^64^I^65^K^66^A, I^65^W^72^A and E^83^S^84^D^85^R^86^G^87^R^88^A, involved residues oriented away from A28, implying a different interfering mechanism from the mutations in the H2 ectodomain at the C-terminus. Accordingly, we speculate that these residues may interact with other EFC components or viral membrane lipids such that alanine mutation abolishes the local protein-protein and/or protein-lipid interactions. This hypothesis is supported by our co-IP results presented in Figure 6. Nevertheless, further work is needed to clarify the role of this long helix (aa51-90) in EFC formation and its mechanistic importance to EFC-induced membrane fusion.

To explore the molecular basis of EFC-mediated membrane fusion, aside from determining the entire EFC complex structure, it is also crucial to decipher the structure and function of individual components to underpin the mechanism of how this eleven-protein machinery functions. In the current study, we obtained a crystal structure of the vaccinia virus H2 ectodomain (i.e., tH2). Through biological and biochemical analyses, we have identified surface loops on the H2 C-terminus crucial for viral infectivity, interaction with A28, and MV-triggered fusion activity. Finally, although the structure of the H2-A28 subcomplex remains to be deciphered, together with the findings from a concurrent study on vaccinia A28 protein [51], our H2 structure enables visualization of the binding surface between H2 and A28, thereby enhancing our knowledge of EFC assembly and paving the way toward understanding the molecular mechanism underlying EFC-mediated membrane fusion.

## Acknowledgments

We thank Shiu-Ping Li at the IMB Imaging Core for assistance and B. Moss for providing the vH2i virus. We also thank the staff of beamline BL15A at the National Synchrotron Radiation Research Center in Hsinchu (NSRRC) in Hsinchu, Taiwan, for their help in X-ray crystal data collection. The work is supported by grants from Academia Sinica (AS-IDR-112-02 to W.C.) and the National Science and Technology Council (110-2320-B-001-015-MY3 to W.C.).

